# Excess of Deleterious Mutations around HLA Genes Reveals Evolutionary Cost of Balancing Selection

**DOI:** 10.1101/053793

**Authors:** Tobias L. Lenz, Victor Spirin, Daniel M. Jordan, Shamil R. Sunyaev

**Affiliations:** Division of Genetics, Department of Medicine, Brigham and Women’s Hospital, Harvard Medical School, Boston, MA 02115, USA; Evolutionary Immunogenomics, Department of Evolutionary Ecology, Max Planck Institute for Evolutionary Biology, 24306 Plön, Germany; DProgram in Medical and Population Genetics, The Broad Institute, Cambridge, MA 02142, USA

**Keywords:** balancing selection, exome, simulations, deleterious variation, mutation load, MHC/HLA

## Abstract

Deleterious mutations are expected to evolve under negative selection and are usually purged from the population. However, deleterious alleles segregate in the human population and some disease-associated variants are maintained at considerable frequencies. Here we test the hypothesis that balancing selection may counteract purifying selection in neighboring regions and thus maintain deleterious variants at higher frequency than expected from their detrimental fitness effect. We first show in realistic simulations that balancing selection reduces the density of polymorphic sites surrounding a locus under balancing selection, but at the same time markedly increases the population frequency of the remaining variants, including even substantially deleterious alleles. To test the predictions of our simulations empirically, we then use whole exome sequencing data from 6,500 human individuals and focus on the most established example for balancing selection in the human genome, the major histocompatibility complex (MHC). Our analysis shows an elevated frequency of putatively deleterious coding variants in non-HLA genes localized in the MHC region. The mean frequency of these variants declined with physical distance from the classical HLA genes, indicating dependency on genetic linkage. These results reveal an indirect cost of the genetic diversity maintained by balancing selection, which has hitherto been perceived as mostly advantageous, and have implications both for the evolution of recombination and also for the epidemiology of various MHC-associated diseases.

## Introduction

A large number of population genetics studies point to the existence of numerous mildly deleterious alleles segregating in the human population (Henn et al. 2015). This abundance of deleterious alleles is evident from the comparison of human DNA polymorphism to human-chimpanzee sequence divergence (Bustamante et al. 2005), from the analysis of allele frequency distribution (Kryukov et al. 2007; Boyko et al. 2008; Kryukov et al. 2009) and estimated allelic ages (Kiezun et al. 2013). Deleterious alleles are present in individual human genomes at functionally significant sites in both protein coding genes and regulatory regions (Maurano et al. 2012; Fu et al. 2013). Understanding the maintenance of deleterious variation in the population is critically important for evolutionary models of complex traits including common human diseases. Persistence of common diseases is paradoxical from the evolutionary standpoint, and the stable existence of deleterious variation in spite of the action of purifying selection requires an explanation.

Different hypotheses have been put forward to explain the occurrence of deleterious genetic variants in the population. Mutation-selection balance in combination with demography and genetic drift certainly account for a significant part of the genetic load and for the persistence of rare deleterious variation (Dudley et al. 2012; Fu et al. 2013). The occurrence of more frequent variants, on the other hand, is commonly hypothesized to involve selective forces favoring genetic diversity. However, selection should not necessarily target the variants in question. With the improved understanding of the recombination landscape and of the extent of genetic linkage, it is becoming increasingly clear that many sites in the genome do not evolve completely independently and allele frequency changes may be influenced by variation at neighboring linked sites (Gillespie 2000; Charlesworth et al. 2003; Barton 2010).

Previous research has already addressed the evolutionary fate of genetic variation around sites under selection, focusing mostly on neutral variation. Background selection has been shown to result in the reduction of neutral diversity surrounding regions under purifying selection (Charlesworth et al. 1993). Genetic hitchhiking events associated with positive selection generally eliminate surrounding genetic variation. On the other hand, they transiently increase allele frequencies of variants residing on the same haplotype as the positively selected variant (Maynard Smith and Haigh 1974; Fay and Wu 2000; Chun and Fay 2011), and theoretical analysis and empirical data suggest that hitchhiking events can also elevate the frequency of deleterious variants (Chun and Fay 2011; Hartfield and Otto 2011; Marsden et al. 2016). Conversely, existing recessive deleterious variation may also slow the fixation of beneficial alleles on the same haplotype (Assaf et al. 2015). However, since hitchhiking eventually reduces the absolute amount of genetic diversity, it is not sufficient to explain regions with generally elevated levels of sequence diversity across the genome.

In contrast, balancing selection leads to a long-term persistence of common genetic variation in surrounding loci (Charlesworth 2006; Gao et al. 2015). Consequently, it has been proposed that balancing selection may also lead to an excess of deleterious variation around the balanced locus. This scenario is supported by studies of neutral variation that showed increased variation in regions linked to loci under both simple and multi-locus balancing selection (Kaplan et al. 1988; Grimsley et al. 1998; O'hUigin et al. 2000; Navarro and Barton 2002). However, a comprehensive theoretical and empirical investigation of this scenario for deleterious variants, which is of critical interest for both population genetics and medical genetics, is lacking so far.

The prevalence of balancing selection in the human population and its role in shaping genetic variation is still debated. In contrast to models developed in the 1960s (e.g Lewontin and Kojima 1960), studies of the genomic era have generally assumed that balancing selection is an exception rather than the rule (Asthana et al. 2005; Bubb et al. 2006). However, an accumulating number of recent studies are identifing genomic features characteristic of balancing selection (Bustamante et al. 2005; Andrés et al. 2009; Fumagalli et al. 2009; Sellis et al. 2011; Gokcumen et al. 2013; Leffler et al. 2013; DeGiorgio et al. 2014; Teixeira et al. 2015).

A classical locus to investigate the effect of balancing selection on multiple sites and its influence on neighboring variation is the major histocompatibility complex (MHC). The MHC is one of the most prominent examples of balancing selection in the vertebrate genome. This is thought to be due to the so-called classical MHC genes (in humans called Human Leukocyte Antigen; HLA), whose products present antigenic peptides on the cell surface and by that play a key role in the adaptive immune response (Neefjes et al. 2011). These classical HLA genes are scattered across the MHC region and exhibit exceptional allelic polymorphism and an extreme level of heterozygosity, which is thought to increase pathogen resistance and thus to be maintained by pathogen-mediated balancing selection (Trowsdale 2011). Based on well-documented signatures of balancing selection at the classical HLA genes, this gene complex thus provides a perfect study system to explore the effect of balancing selection on the frequency of linked deleterious variation. In fact, a striking feature of the MHC, potentially related to the maintenance of deleterious variation, is that the MHC region is highly enriched in variants associated with common human diseases identified by Genome-Wide Association Studies (GWAS; fig. 1). Many of the GWAS peaks are not caused by variation in classical HLA genes, and a significant number of phenotypes associated with genetic variants within the MHC region are not classic autoimmune or infectious diseases (Trowsdale 2011).

**Figure 1:**
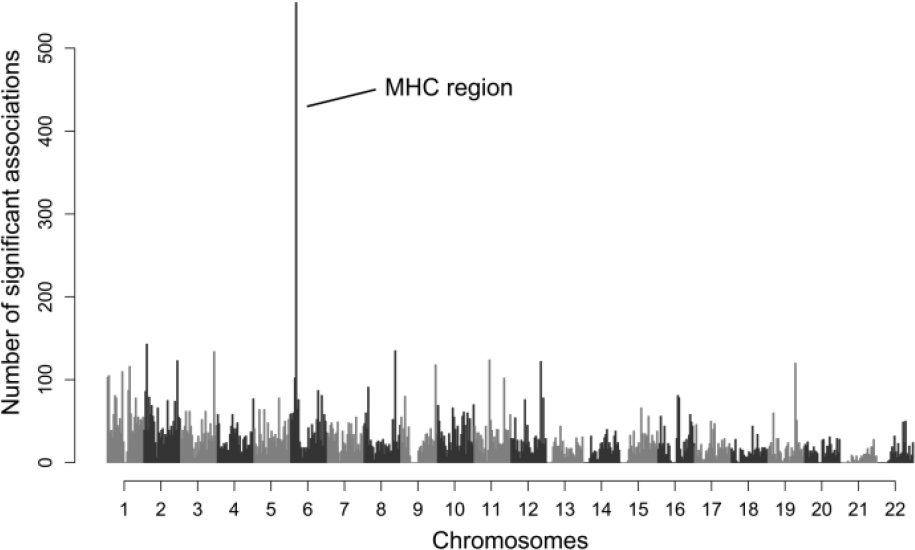
**Number of significant GWAS associations along the genome.** The chromosomal location of significant trait associations from genome-wide association studies (GWAS, N=18,682) are shown for all autosomes. Data from NHGRI GWAS catalog.

To investigate the potential of balancing selection for maintaining deleterious genetic variation, we first use a forward simulation approach and then analyze empirical data to test the predictions of the simulations. Previously, the large scale sequencing data required to compare the mutational load in the MHC region against the rest of the genome has been lacking. Here we make use of a whole exome sequence data set of about 6,500 individuals, which provides detailed population-level information of coding sequence variation across the entire human genome.

## Results

### Simulating Multi-Locus Balancing Selection

We used forward simulations to investigate the effect of multi-locus balancing selection at HLA genes on the evolution of deleterious genetic variants throughout the MHC region. The simulations were based on empirical parameter values from the human MHC region and included a central HLA gene, surrounded by non-HLA regions that represented the variation in neighboring non-HLA genes. In order to simulate multi-locus selection, multiple scattered sites along the virtual HLA gene evolved under different HLA selection scenarios: either neutrally, under balancing selection, or, as a contrast, under recurrent sweeps of positive selection. We then explored the effect of these different HLA selection scenarios on the evolution of variants in HLA-neighboring regions that evolved themselves under different levels of purifying selection.

Our simulations showed that multi-locus balancing selection at HLA generally reduces the number of segregating sites in the neighboring sequence regions compared to a scenario with no selection at HLA (fig. 2a). With moderate or high levels of purifying selection acting at neighboring sites, the number of segregating sites was even lower. Interestingly, the reduction in the density of segregating sites by balancing selection was mainly due to the removal of sites with rare variants (fig. 2b). And as nucleotide diversity (*π*) is strongly dependent on variant frequencies, the removal of rare variants had little effect on overall diversity in the neighboring regions. Instead, balancing selection at the HLA led to a significant increase in nucleotide diversity in the neighboring regions surrounding the HLA and was elevated even in the presence of moderate purifying selection (fig. 2c). Nucleotide diversity increased in spite of the reduction in number of segregating sites because derived allele frequencies at the remaining sites were, on average, strongly elevated, resulting in a site frequency spectrum that was enriched with intermediate frequency alleles (fig. 3a). This increase in the number of segregating sites with intermediate frequency alleles then caused an overall increase in nucleotide diversity, irrespective of the significant loss of rare variants around the HLA gene. As expected, increasing purifying selection in the surrounding region reduced allele frequencies overall. However, relative to the other HLA selection scenarios, balancing selection at the HLA in all simulated cases maintained a significant fraction of deleterious variation at moderate frequencies counteracting the effect of purifying selection (Wilcoxon rank-sum test, all *P* < 0.001; fig. 3b–fig. 3d). The effect of balancing selection on variant frequency in the neighboring regions was strongly dependent on the rate of recombination: Artificially increasing the recombination rate by an order of magnitude basically removed the effect, while an equivalent reduction in recombination led to a substantially stronger enrichment for intermediate frequency variants at all levels of purifying selection (**supplementary figs. S1 & S2**, Supplementary Material online).

**Figure 2:**
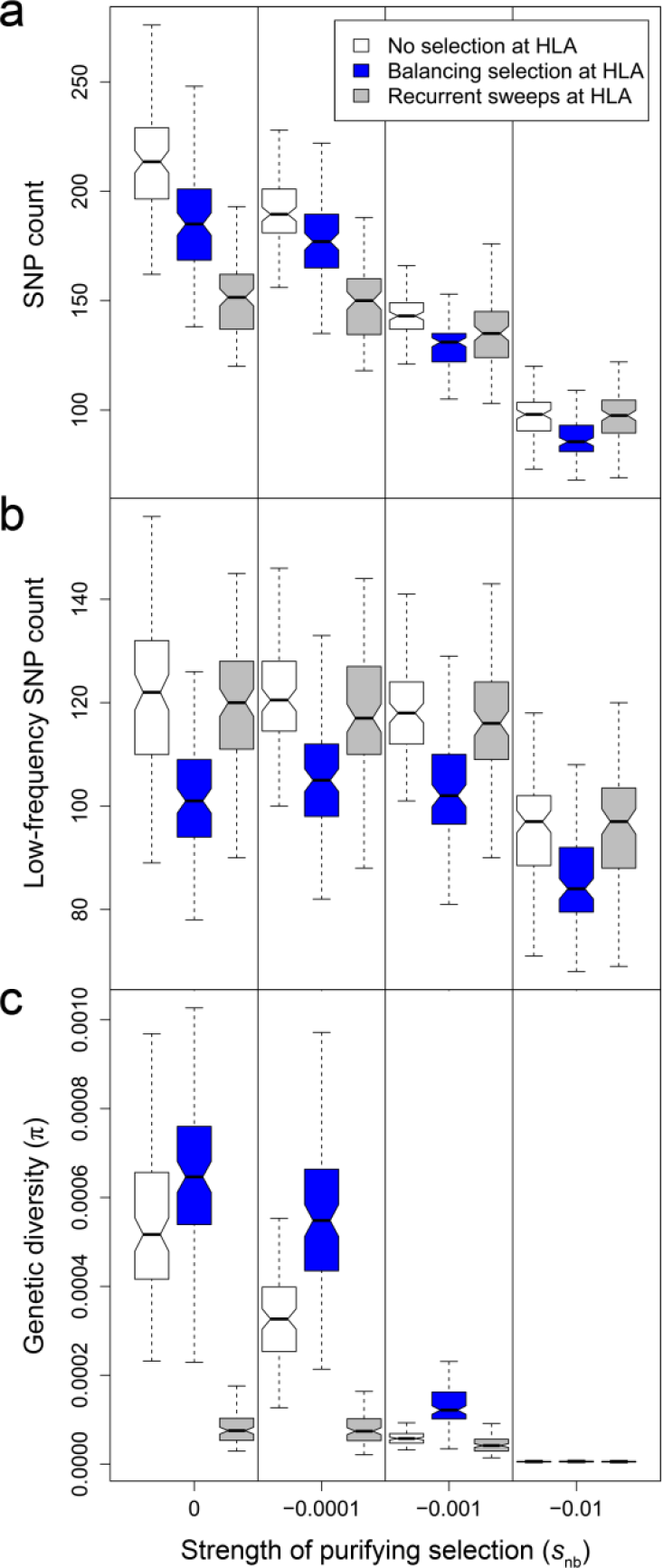
**Simulated polymorphism in regions surrounding the HLA gene under different selection scenarios.** Polymorphic sites (SNPs) along the regions around an HLA gene are derived from simulations with three different selection scenarios on the HLA gene (white: no selection on HLA, blue: balancing selection, grey: recurrent sweeps of positive selection). Standard box plots show the median number of (**a**) all SNPs, (**b**) only SNPs with derived allele frequency < 0.01, and **(c)** the genetic diversity (π) across all sites. Variants in surrounding regions evolved neutrally (S_nb_ = 0) or under co-dominant purifying selection with S_nb_ = −0.0001, S_nb_ = −0.001, or S_nb_ = −0.01, respectively. Nonoverlapping notches between box plots indicate significant difference. Note the different y-axis scales.

**Figure 3:**
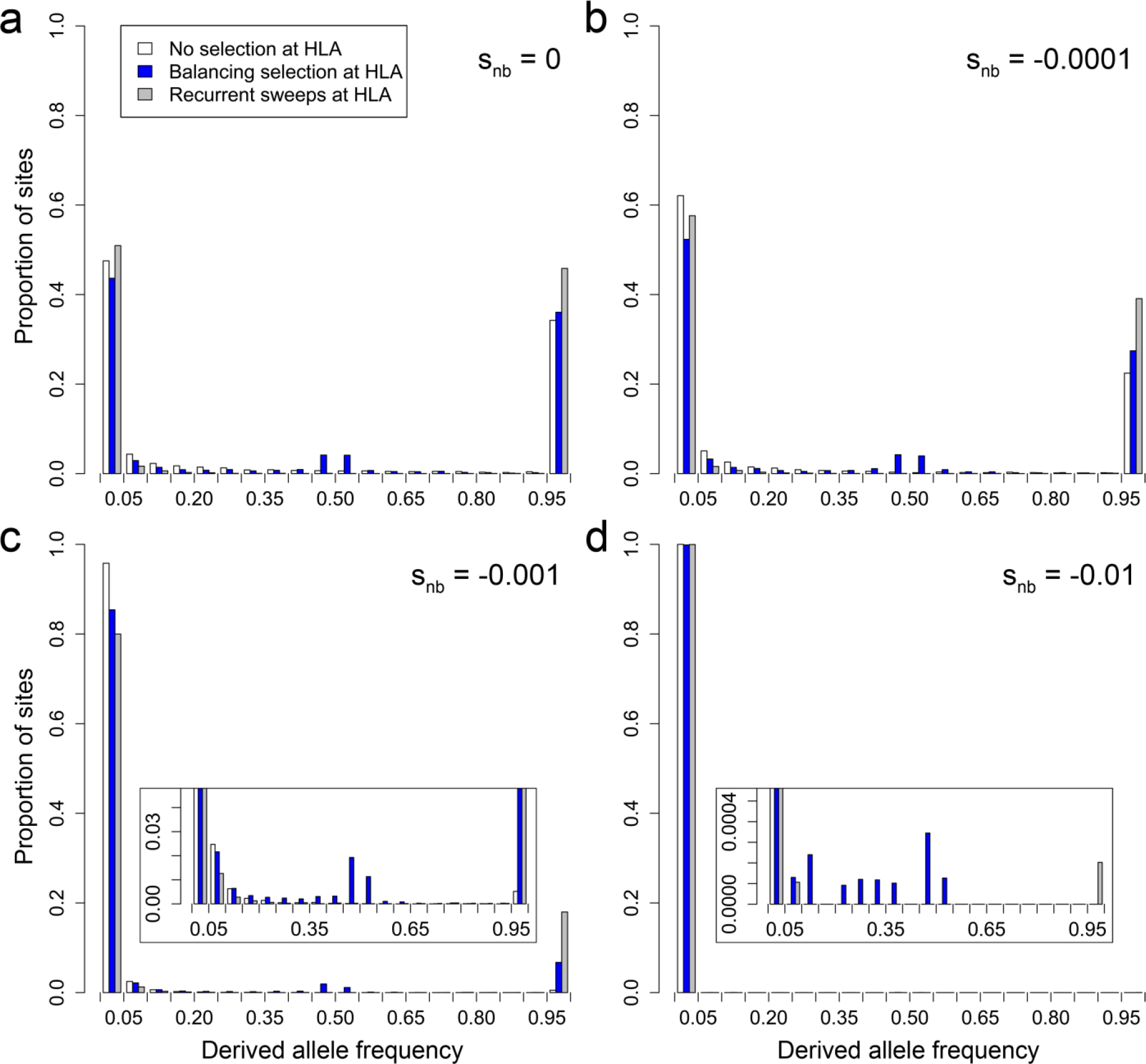
**Simulated site frequency spectrum of variants surrounding the HLA gene under different selection scenarios.** Derived allele frequencies along the regions around an HLA gene are derived from simulations with three different selection scenarios on the HLA gene (white: no selection on HLA, blue: balancing selection, grey: recurrent sweeps of positive selection). Variants in neighboring regions evolved (**a**) neutrally (S_nb_ = 0) or under co-dominant purifying selection with (**b**) S_nb_ = −0.0001, (**c**) S_nb_ = −0.001, or (**d**) S_nb_ = −0.01, respectively. Note the different y-axis scales in the zoomed insets of panels (**c**) and (**d**) for better visualization.

We contrasted these results with the alternative scenario of recurrent complete selective sweeps at the HLA due to continuous environmental change, e.g. constantly adapting or newly arising pathogens with ‘sweeping’ selective effects (e.g. the Bubonic plague). Similarly to the balancing selection scenario, recurrent sweeps in HLA genes led to a reduction in the number of segregating sites in the neighboring regions (fig. 2a). Compared to the balancing selection scenario, this reduction in SNP density is stronger if there is no purifying selection on the surrounding regions and weaker when purifying selection exerts pressure on the neighboring sequence. This reduction is primarily affecting SNPs with common derived alleles, with almost no removal of rare variants (fig. 2b). Consequently, in sharp contrast with the balancing selection scenario, nucleotide diversity is reduced rather than elevated (fig. 2c).

Overall, our simulations showed that multi-locus balancing selection can lead to a distinct signature of a reduced number of polymorphic sites with elevated allele frequencies. The elevated site frequency spectrum suggests that balancing selection has the realistic potential to significantly increase of even frequencies of strongly deleterious variants in regions around the classical HLA genes.

### Exome Sequencing Data

We used SNP variation data from the whole-exome sequence dataset of the NHLBI GO Exome Sequencing Project (from a total of about 6,500 humans and 17,684 genes, including 124 genes within the MHC region). Excluding eleven of the latter that are classical HLA genes, expected to evolve under balancing selection, our analyses of the MHC region included variant data from 113 genes. The median number of polymorphic sites averaged 48 for these genes, significantly lower than the exome-wide median of 74 (Wilcoxon rank-sum test, *P* < 0.001; fig. 4a) and was mainly due to a difference in the number of sites with low frequency variants (fig. 4b). Our simulations also showed fewer polymorphic sites under the balancing selection scenarios than without it. The reduced number of polymorphic sites affected all SNP categories, including variants likely to be deleterious (as predicted by PolyPhen-2 (Adzhubey et al. 2010); this distinguishes the effect of balancing selection from other scenarios, such as genetic hitchhiking during selective sweeps where the higher frequency synonymous variants are preferentially removed from the population (Chun and Fay 2011). In contrast, the population frequency of derived alleles at polymorphic sites was substantially elevated within the MHC region compared to the rest of the genome (fig. 5a).

**Figure 4:**
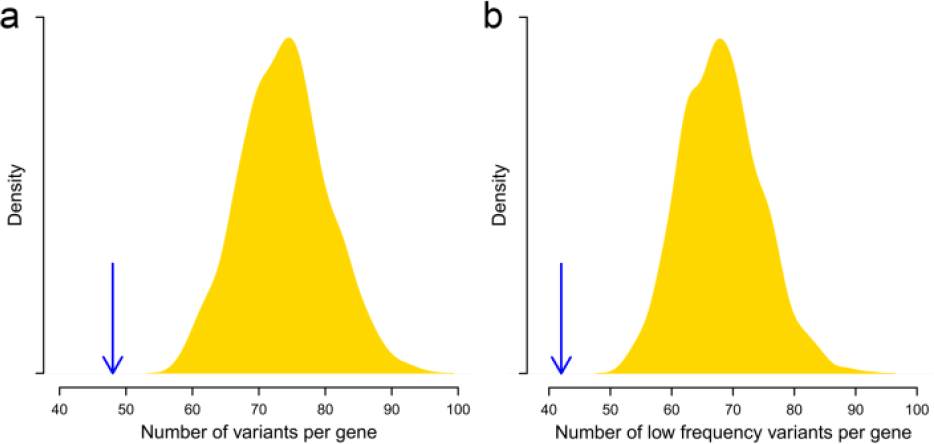
**Observed average number of polymorphic sites per gene.** The median number of polymorphic sites per gene is averaged over the 113 genes represented in the MHC region (blue arrow, excluding classical HLA genes). Also shown is the distribution of equivalent values for 1,000 Monte-Carlo-sampled sets of 113 random genes from the entire exome. Represented are (**a**) all SNPs and (**b**) only SNPs with derived allele frequency < 0.01.

**Figure 5:**
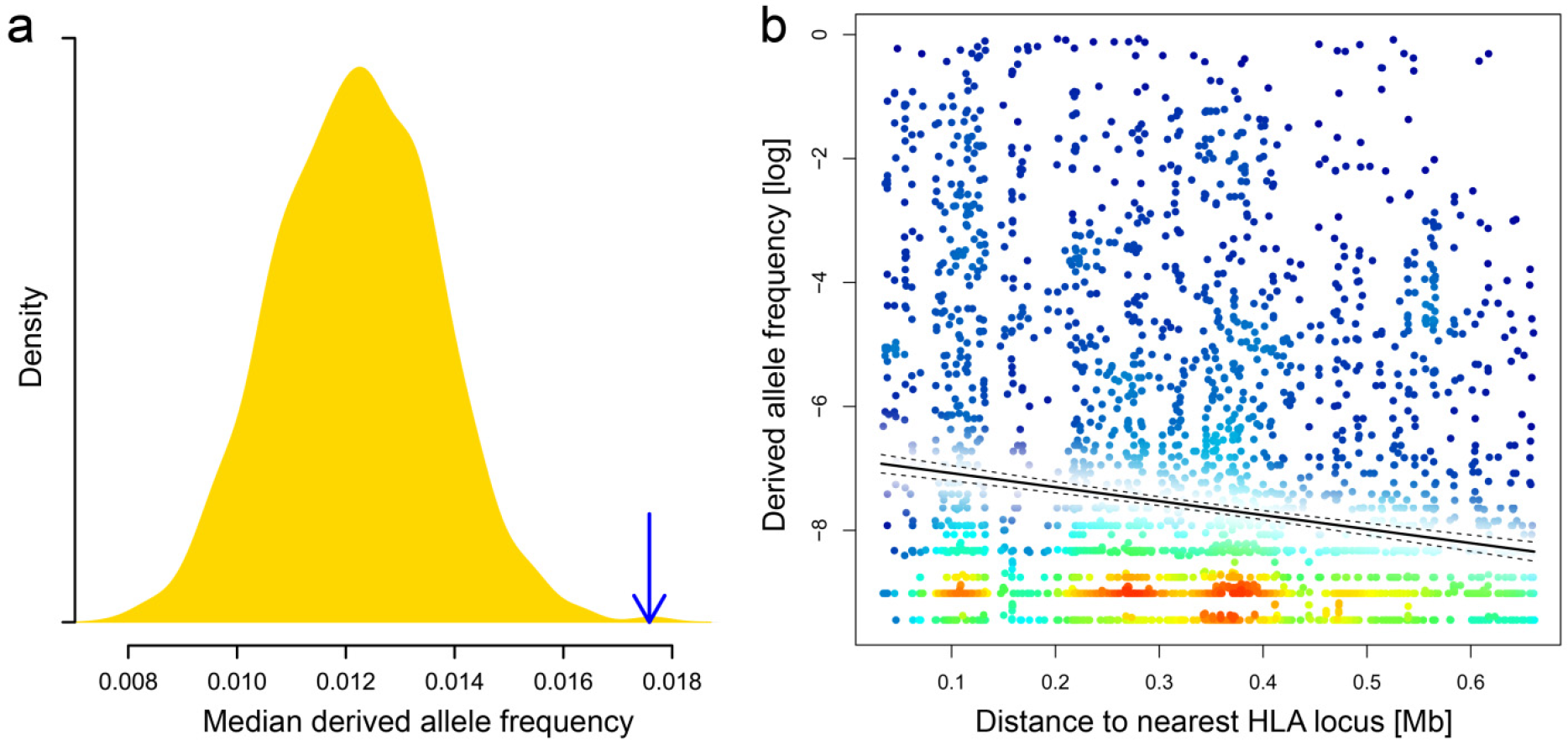
**Observed distribution of derived allele frequencies.** Derived alleles are defined as the non-reference allele at polymorphic sites in the Exome Sequencing Project data. **(a)** The median frequency of derived alleles per gene is averaged over the 113 genes represented in the MHC region (blue arrow, excluding classical HLA genes). The yellow density curve shows the distribution of equivalent values for 1000 Monte-Carlo-sampled sets of 113 random genes from the entire exome. **(b)** Derived allele frequency at polymorphic sites (N = 4,175) in the MHC region is shown in relation to distance to the nearest classical HLA locus. Solid and dashed lines indicate linear fit and 95% confidence intervals, respectively. Color shading indicates local data point density for improved visualization of the extent of overlapping points in the plot, ranging from blue (low density) via green and yellow to red (high density).

This result is not caused by local differences in mutation rate, since the rate of *de novo* mutations (as estimated from trio sequencing; Francioli et al. 2015) in the MHC region does not differ from the average exome-wide rate (Wilcoxon rank-sum test, MHC vs. whole exome, *P* > 0.05). The frequencies of derived alleles correlated with their position relative to the nearest classical HLA locus: derived alleles at sites physically close to an HLA locus were at higher frequencies than at sites further away (Spearman's ρ = −0.14, *P* < 0.001; fig. 5b), suggesting that their frequencies are influenced by linkage disequilibrium to HLA genes.

Since we were especially interested in potentially deleterious variants, we focused our analyses on loss-of-function variants and missense variants classified as “probably damaging” by PolyPhen-2 (Adzhubey et al. 2010), which are expected to evolve under moderate to strong purifying selection. This expectation was supported by a substantially lower exome-wide derived allele frequency of “probably damaging” variants, compared to variants classified as “benign” (Wilcoxon rank-sum test, both P < 0.001; **supplementary fig. S3**, Supplementary Material online). We found a significant elevation in average derived allele frequencies within the MHC region, for both probably damaging missense variants and loss-of-function variants, compared to the average frequency across the entire exome (Wilcoxon rank-sum test, both *P* < 0.001; fig. 6). This shift in the variant frequencies was observed within both African American and European American subpopulations of the ESP6500 data (**supplementary note S1, supplementary table S1 and supplementary fig. S4**, Supplementary Material online).

**Figure 6:**
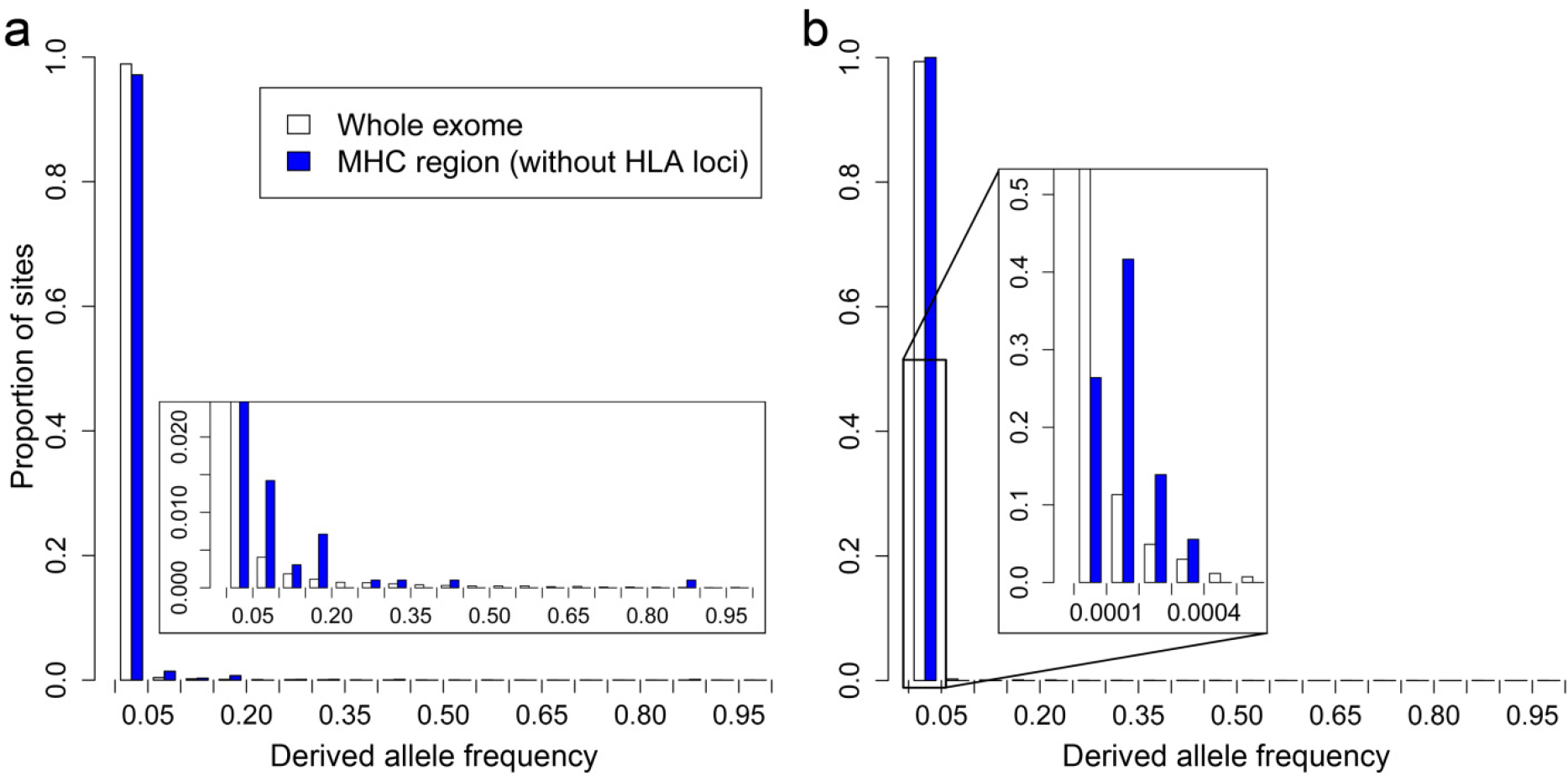
**Observed site frequency spectrum of deleterious variants.** The site frequency spectra of (**a**) probably damaging and (**b**) loss-of-function variants are shown for the entire exome (white bars) and the MHC region only (blue bars, excluding the classical HLA loci).

Overall, we identified a substantial number of non-HLA genes throughout the MHC region, carrying deleterious variants at high frequencies (**supplementary table S2**, Supplementary Material online). For some of these genes, the average frequency of probably damaging variants was more than two orders of magnitude above the genome-wide average. Such genes include *MICA* (6 variants predicted to be ‘probably damaging’ with mean derived allele frequency, or DAF, of 9.7%), *PSORS1C1* (2 variants with DAF of 8.8%), and *CFB* (9 variants with DAF of 2.0%). **Supplementary table S2** (Supplementary Material online) lists potential diseases reported in connection with these genes, including common autoimmune diseases (e.g. psoriasis, rheumatoid arthritis), cancer (e.g. prostate cancer, hepatocellular carcinoma), and mental disorders (e.g. Alzheimer's disease, schizophrenia).

## Discussion

Our simulations of genomic regions around a classical HLA gene showed that balancing selection has a similar effect on the number of linked variants as recurrent sweeps of positive selection in the way it reduces the absolute number of segregating sites. However, while positive selection simultaneously reduces the frequency of variants at the remaining sites, balancing selection has the opposite effect and substantially increases derived allele frequencies, leading to a site frequency spectrum with an excess of intermediate frequency alleles. As expected, derived allele frequencies also depended on the strength of selection against the deleterious variants. This is in agreement with previous studies on the evolution of neutral variation around balanced loci, showing generally elevated sequence diversity (Kaplan et al. 1988; Grimsley et al. 1998; Horton et al. 1998; O'hUigin et al. 2000; Navarro and Barton 2002; Connallon and Clark 2013).

Results from deep population sequencing data supported our simulation results empirically, by showing the same pattern of a reduced number of polymorphic sites but elevated allele frequencies in the MHC region compared to the rest of the exome. This observation held for both loss-of-function mutations, which are likely to be highly deleterious, as well as presumably deleterious variants, some of which reside in genes with known disease associations, again suggesting moderate to strong purifying selection. The dependency of allele frequencies on the proximity to classical HLA genes supports the notion that balancing selection acts on neighboring variation via linkage disequilibrium. Independent support for our findings comes also from two earlier studies that described elevated levels of genetic diversity around specific MHC genes and suggested that this might be maintained by balancing selection on the given neighboring MHC locus (Horton et al. 1998; Shiina et al. 2006).

The observed reduction in the number of polymorphic sites in regions close to loci under multi-locus balancing selection both in simulations and empirical data is an interesting observation, given the expectation that balancing selection generally increases genetic diversity around the loci under selection (Navarro and Barton 2002; Charlesworth 2006). This is also a significant difference to the effect of genetic hitchhiking after selective sweeps, which has been shown to reduce the number of neutral variants in neighboring regions but not to affect the number of deleterious variants (Chun and Fay 2011). However, the unexpected reduction in the number of polymorphic sites with balancing selection at the neighboring HLA is likely due to the recurrent occurrence of new mutations at one of the balanced HLA sites, a central aspect of our multi-locus simulation model. Such a new variant, being subject to overdominant selection, would drive the haplotype on which it occurred to higher frequency (but not to fixation), dragging along any linked neighboring variant. At the same time the rising frequency of this haplotype would replace other haplotypes that don’t carry the new balanced variant. Consequently, a polymorphic site linked to a balanced locus can have two fates: Either one of its minor alleles is present on the new balanced haplotype, so that its frequency rises together with the balanced haplotype, or the new balanced haplotype does not carry the site’s minor alleles, so that its frequency declines. For very low frequency variants (e.g. singletons) the latter fate will potentially lead to a complete loss of the minor allele at this neighboring site, and thus to a complete loss of the segregating site. Together these two mechanisms lead to the signature that we observe: Less polymorphic sites around the HLA gene but a higher derived allele frequency at those sites that remain in the population.

Such a scenario of recurrent but incomplete sweeps could indeed result from several mechanisms of balancing selection, such as negative frequency-dependent selection and fluctuating selection in time and space, which are also thought to affect the HLA (Meyer and Thomson 2001; Spurgin and Richardson 2010). Previous theoretical work has even suggested that symmetrical overdominance alone is not sufficient to explain the entire allelic diversity seen at the HLA (De Boer et al. 2004). Furthermore, the HLA exhibits identity-by-descent patterns that cannot be explained by ancient balancing selection, suggesting ongoing selection and very recent selective events (Albrechtsen et al. 2010), for instance through local adaptation after human migration and/or drastic epidemic events in recent human history (Zhou et al. 2016). All these processes are likely to contribute to the distribution of linked deleterious variation around the classical HLA genes.

Mechanistically, it had also been proposed that the excessive heterozygosity in the MHC region, caused by balancing selection, may reduce the phenotypic expression of recessive deleterious variants, rendering purging mechanisms less effective and thus allowing for the accumulation of a recessive 'sheltered load' that could ultimately even contribute to inbreeding depression (Charlesworth and Willis 2009). Such a scenario was first invoked to explain the unexpectedly long terminal branches in genealogies of genes under balancing selection (Uyenoyama 1997; for a schematic see van Oosterhout 2009) and has since been investigated by theoretical and empirical studies on the self-incompatibility determining S locus, the most compelling example for balancing selection in plants. Those studies suggested that recessive deleterious mutations may accumulate around the S locus, presumably protected from purifying selection by the excessive heterozygosity in this region, and, once established, contribute to the disadvantage of homozygotes (Stone 2004; Llaurens et al. 2009). Such a 'sheltered load' mechanism has later also been proposed to contribute to the diversity found in the MHC region in vertebrates (van Oosterhout 2009; Llaurens et al. 2012). However, in contrast to previous work, our models are based on empirical parameter values, showing an accumulating deleterious load under realistic levels of recombination. In addition we here provide direct empirical evidence of elevated deleterious variation in the human MHC region. While it is conceivable that some of these observed deleterious variants represent such a 'sheltered load', our simulations assume additive (and not recessive) effects for all non-HLA variants, indicating that there must be other mechanisms besides the ‘heterozygosity shelter’ for recessive variants that contribute to the observed deleterious load.

Contributing to the observed excess of deleterious variants could also be a local alteration of the effective population size, and consequently of the efficacy of selection. Balancing selection, and specifically symmetrical overdominance, can lead to a limited number of dominating haplotypes that are stably balanced at intermediate frequencies in the population. Without recombination, such dominating haplotypes could be seen as essentially subdividing the population in this genomic region (Charlesworth 2006), so that genetic drift of linked variants occurs only within the same haplotype background. This would result in a locally reduced effective population size per haplotype, rendering selection less efficient within a given haplotype background, even though, when estimated across all haplotypes, balancing selection increases the effective size of the total population (Charlesworth 2009). This phenomenon is strongly based on genetic linkage, and with recombination the effect on linked variants will quickly break down with increasing distance to the balanced locus (Hudson and Kaplan 1988). Such a dependence between the excess deleterious load and the rate of recombination could indeed be observed both in the simulations and in the empirical data (here using physical distance along the chromosome as a proxy for genetic linkage).

Overall, our results show that balancing selection has the potential to maintain deleterious genetic variants at considerable frequencies in natural populations, especially if they are only moderately deleterious, e.g. causing late-onset autoimmunity. The extent of the deleterious mutational load on a given MHC haplotype will ultimately depend on its aggregate detrimental effect and the role of linked HLA alleles, resulting from an evolutionary trade-off between the antagonistic effects of the number (and severity) of accumulating deleterious variants and the selective advantage of individual allelic variability at the classical HLA genes. Interestingly, more divergent HLA genotypes are assumed to confer broader immune-surveilance against antigens (Lenz 2011), and a positive correlation between the pathogen diversity in a given population and its HLA allele pool diversity has been reported (Prugnolle et al. 2005). It could thus be hypothesized that environments with higher pathogen diversity, and thus stronger selection for high individual allelic diversity at HLA, may allow for the maintenance of MHC haplotypes with a larger deleterious load (Dean et al. 2002). Once the selective pathogenic pressure declines (either through host migration to a less pathogenic environment or through environmental changes), this could then result in increased expression of phenotypes associated with the deleterious load. Indeed, first evidence associating heterozygosity at the HLA with increased risk for some autoimmune diseases has been reported (Lenz et al. 2015), but further analyses are required to evaluate this hypothesis.

The observed effect of balancing selection on linked variation is evidently dependent on recombination rate around the locus under balancing selection (Hudson and Kaplan 1988; Barton 2010). The MHC region is in fact known to harbor a substantial number of recombination hot-spots (de Bakker et al. 2006), which might have evolved due to the above described dynamics. On the other hand, the MHC region also exhibits even more extreme cases of multi-locus balancing selection than modeled here, where LD spans across separate HLA loci. For instance, the well-documented COX haplotype covers the entire MHC region and has a population frequency of about 10% in Northern Europe (Stewart et al. 2004). Epistasis among different HLA genes, for instance due to advantageous allele combinations, may select against recombination and maintain such long-range haplotypes (Penman et al. 2013). The genome-wide extent of the observed effect remains to be investigated, but given the increasing number of regions detected to evolve under balancing selection (Leffler et al. 2013), there is reason to expect that this mechanism may significantly contribute to the prevalence of heritable human diseases.

## Materials and Methods

### GWAS Summary

Data for GWAS hit summary was downloaded from the NHGRI GWAS catalog (available at: http://www.genome.gov/gwastudies, accessed [02/15/2015]) (Welter et al. 2014). This catalog represents a comprehensive collection of published genome-wide association studies (currently 1,755 studies), including the investigated trait and the chromosomal location of one or more (median: 4) independent associations per study. For some traits the data contains multiple studies (top five: type 2 diabetes: 37, breast cancer: 26, schizophrenia: 22, body mass index: 21, height: 21; median number of studies per trait: 1), but preferentially lists novel associations to avoid redundancy. While this data may suffer from a bias towards over-studied traits, it also reflects the frequency and thus importance of a given trait in the population. We focused only on autosomal associations and chromosome length was standardized to 30 location bins per chromosome for better visualization.

### Simulating Multi-Locus Balancing Selection

In order to test whether balancing selection on a particular locus can lead to a shift in the site frequency spectrum of adjacent genomic regions and prevent deleterious mutations from being purged by purifying selection, we employed a forward simulation approach. To this end, we developed the program Forward Simulation (available at http://forwardsimulation.sourceforge.net), which is based on the Wright-Fisher model and allows to define distinct selection regimes for separate regions or sites of the simulated genome. The program uses a multiplicative fitness framework with diallelic sites, with fitness *f* = (1+s*h*)*^n^*, where s and *h* are the selection and dominance coefficients per site, respectively, and *n* is the number of sites in the genome. A negative selection coefficient s leads to negative (purifying) selection and vice versa. The simulated genome contained a virtual HLA gene of the median genomic length of classical HLA genes (start of first exon to end of last exon, including introns; 5,385 bp). This HLA gene was flanked on both sides by surrounding regions (set to evolve under different levels of purifying selection). The length of these two regions equaled half the median distance between adjacent classical HLA genes in the MHC region (total length: 37,122 bp). Following the multi-locus balancing selection scenario by Navarro and Barton (2002), a number of polymorphic sites along the virtual HLA gene were specified to evolve under the HLA selection regime at the beginning of each simulation (after burnin). Once the sites had been specified, they were free to maintain, lose, or regain polymorphism throughout the simulation. These sites mimic the functional polymorphism seen in classical HLA genes, where sites along the exons coding for the antigen binding groove of the HLA molecule show variation that is thought to evolve under balancing selection (Reche and Reinherz 2003; Furlong and Yang 2008). See **supplementary fig. S5** (Supplementary Material online) for a schematic of the simulated genome. For exploring the effect of balancing selection on surrounding regions in a near-realistic scenario of MHC evolution, we used empirical values from the literature for all fixed parameters: effective population size N_e_ = 10,000 (Takahata et al. 1995), mutation rate p = 1.38e-8 (Scally and Durbin 2012), recombination rate r = 0.44 cM/Mb (de Bakker et al. 2006; Taylan and Altiok 2012). Average selection at the classical HLA genes has been estimated as s = 0.013 (Satta et al. 1994; Slatkin and Muirhead 2000; Yasukochi and Satta 2013). Following Navarro and Barton (2002), we simulate multi-locus balancing selection as symmetrical overdominance, setting the selection coefficient *S*HLA for all sites with HLA selection regime to 0.013 for heterozygotes and to 0 for homozygotes. For simulation scenarios with no selection at HLA, *S*_HLA_ was set to 0. The scenarios with recurrent selective sweeps at the HLA gene were simulated by setting *S*_HLA_ = 0.013, with a dominance coefficient *h* = 0.5, and reverting sites with fixed derived allele back to the ancestral allele in order to allow for sweeps to reoccur. The number of sites to evolve under the HLA selection regime was 117, which corresponds to the average number of amino acid residues coding for the peptide binding domain of the classical HLA molecules. All other sites along the entire genome were set to evolve either neutrally (*S* at non-HLA or ‘neighboring’ sites: *S*_nb_ = 0) or under purifying selection, with *S*_nb_ ranging from −0.0001 to −0.01, and dominance *h* = 0.5 (additive). The simulations were run for 150,000 generations, after an initial burnin of 100,000 generations intended to generate an equilibrium level of neutral variation (***π*** *≈* 4Nμ **supplementary fig. S6**, Supplementary Material online). During the burnin period, selection at the HLA gene was turned off (*S*_HLA_ = 0), so that all sites evolved according to the specified level of purifying selection (*S*_nb_ = 0 to −0.0001), including the possibility for fixation of derived alleles. Each simulated scenario was replicated 100 times. Reported results are based only on variants in the surrounding region, thus excluding the virtual HLA gene.

### Exome Sequencing Data and MHC Region

Genome-wide coding sequence variation data of about 6,500 human individuals was obtained from the Exome Variant Server (NHLBI GO Exome Sequencing Project (ESP6500 release), Seattle, WA; http://evs.gs.washington.edu/EVS/; accessed 11/2012) and parsed with custom scripts. Only variants that passed the internal EVS quality filter were recorded and their number and frequencies averaged per gene to account for differences in cds length among genes. Missense variants were further split into functional categories, as determined by Polyphen-2 (Adzhubey et al. 2010). All functional annotations used here are based on the non-reference allele. For simplicity we subsequently call this the derived allele, even though the true ancestral state may not in all cases be reliably resolved. However, our analyses rely only on the functional annotation of a given allele and are independent of the true ancestral/derived state.

The 3.5 Mb region of the classical major histocompatibility complex (MHC) on chromosome 6 was defined following Horton *et al.* (2004), starting with the gene *ZFP57* at position chr6:29640169 and ending with the gene *HCG24* at position chr6:33115544. This excludes the extended MHC regions which contain large numbers of olfactory receptor and histone genes that are likely subject to different types of selection regimes and are thus outside the scope of this study. Within the classical MHC region, the ESP6500 data provided sequence variation data for 124 genes, which corresponds to 73.4% of the known protein-coding genes in this region. This fraction corresponds well with the genome-wide proportion (73.5%) of genes sufficiently covered by the ESP data. Eleven of the 124 genes covered in the MHC region are classical HLA genes (class I: *HLA*-*A*, -*B*, -*C*; *class II*: *HLA-DPA1*, *-DPB1*, *-DQA1*, *-DQA2*, *-DQB1*, *-DRA*, *-DRB1*, *-DRB5*).

Site frequency spectra (SFS), based on derived allele frequencies, were obtained for each of the variant categories across the entire exome as well as for only the MHC region. SFS were compared using the non-parametric Wilcoxon rank-sum test. Within the MHC region, we also calculated for each variant the distance in base pairs from the nearest classical HLA locus and correlated this distance with the derived allele frequency using Spearman correlation. We furthermore compared observed parameter values in the MHC region with genome-wide expectations by Monte Carlo sampling (1,000 replications) from all covered genes in the ESP6500 data. Data analysis was done in R (ver. 3.1.2) (R Development Core Team 2014).

Disease associations for genes in the MHC region with the highest average frequency of potentially deleterious variants were obtained from NHGRI GWAS catalog (see above) and the NIH Genetic Association Database (available at: http://geneticassociationdb.nih.gov, accessed [12/14/2013]) (Becker et al. 2004).

## Acknowledgements

We thank D. Balick for fruitful discussions and comments on a previous version of the manuscript. P. Polak kindly shared gene-wise mutation rate data. We are also grateful to the constructive and thoughtful comments of the editor and three reviewers that helped to improve the manuscript. This work was supported by German Research Foundation (DFG) grants LE 2593/1-1 and LE 2593/2-1 (to T.L.L.), and National Institutes of Health (NIH) grants R01 GM078598 and R01 MH101244 (to S.R.S. and D.M.J.).

